# VExD: A curated resource for human gene expression alterations following viral infection

**DOI:** 10.1101/2021.12.02.470974

**Authors:** Phillip J Dexheimer, Mario Pujato, Krishna Roskin, Matthew T Weirauch

## Abstract

Much of the host antiviral response is mediated through changes to host gene expression levels. Likewise, viruses induce changes to host gene expression levels in order to promote the viral life cycle and evade the host immune system. However, there is no resource that specifically collects human gene expression levels pre- and post-virus infection. Further, public gene expression repositories do not contain enough specialized metadata to easily find relevant experiments. Here, we present the Virus Expression Database (VExD), a freely available web site and database that collects human gene expression datasets in response to viral infection. VExD contains ∼8,000 uniformly processed samples obtained from 289 studies examining 51 distinct human viruses. We show that the VExD processing pipeline preserves known antiviral responses in the form of interferon-stimulated genes. We further show that the datasets collected in VExD can be used to quickly identify supporting data for experiments performed in human cells or model organisms. VExD is freely available at https://vexd.cchmc.org/.

## Introduction

Viral infection can have profound and lasting impacts on human health and society (Vos et al., 2020). Viruses affect host gene expression levels by altering transcriptional mechanisms in infected cells and by initiating the host’s immune response. Numerous transcriptomic studies of human viral infection have been deposited in public databases such as the Gene Expression Omnibus (GEO) (Barrett et al., 2013). However, descriptions of samples and studies in GEO are freeform text and require either manual curation or machine learning to search at a large scale (Z. Wang et al., 2019). This task is particularly complicated in virus-based studies due to the inconsistent nomenclature of viruses in the literature (Gibbs, 2003). As a recent example, the virus that causes COVID-19 was named in the earliest publications as “Wuhan coronavirus” (Liu & Saif, 2020), then “2019-nCov” (Xie & Chen, 2020), before finally settling into the accepted name of “SARS-CoV-2” (Gorbalenya et al., 2020).

In addition to using different nomenclature for the same virus, different studies employ different experimental assays and analysis pipelines to quantify gene expression levels and changes between conditions. These differing conditions make comparison between studies difficult since true biological differences often cannot be distinguished from technical or analytical artifacts.

Collectively, these issues greatly impede progress on studies of the human response to virus infection. To the best of our knowledge, the only resource that attempts to provide human gene expression signatures in response to infection by a variety of viruses is Harmonizome (Rouillard et al., 2016). One of the datasets in that resource (https://maayanlab.cloud/Harmonizome/dataset/GEO+Signatures+of+Differentially+Expressed+Genes+for+Viral+Infections) contains 366 signatures of viral infection. However, these signatures only cover 15 distinct viruses, no attempt was made to standardize virus nomenclature, and only limited information is provided regarding data provenance.

To address the need for uniformly annotated and processed gene expression data from human cells infected with viruses, we present VExD, the Virus Expression Database **(Figure 1)**. VexD contains a manually curated list of 7,903 samples from 289 gene expression studies of infection by 51 distinct viruses. All studies have been subjected to uniform quality control and data processing to maximize comparability across studies. The VExD web interface is easily browsable and searchable and provides automatically generated figures for comparing human gene expression levels across studies and conditions. VExD data are available for download in multiple convenient formats and through an Application Programming Interface (API). VExD is a unique and accessible resource for globally and uniformly assessing the effect of human viruses on gene expression levels.

**Figure 1.**
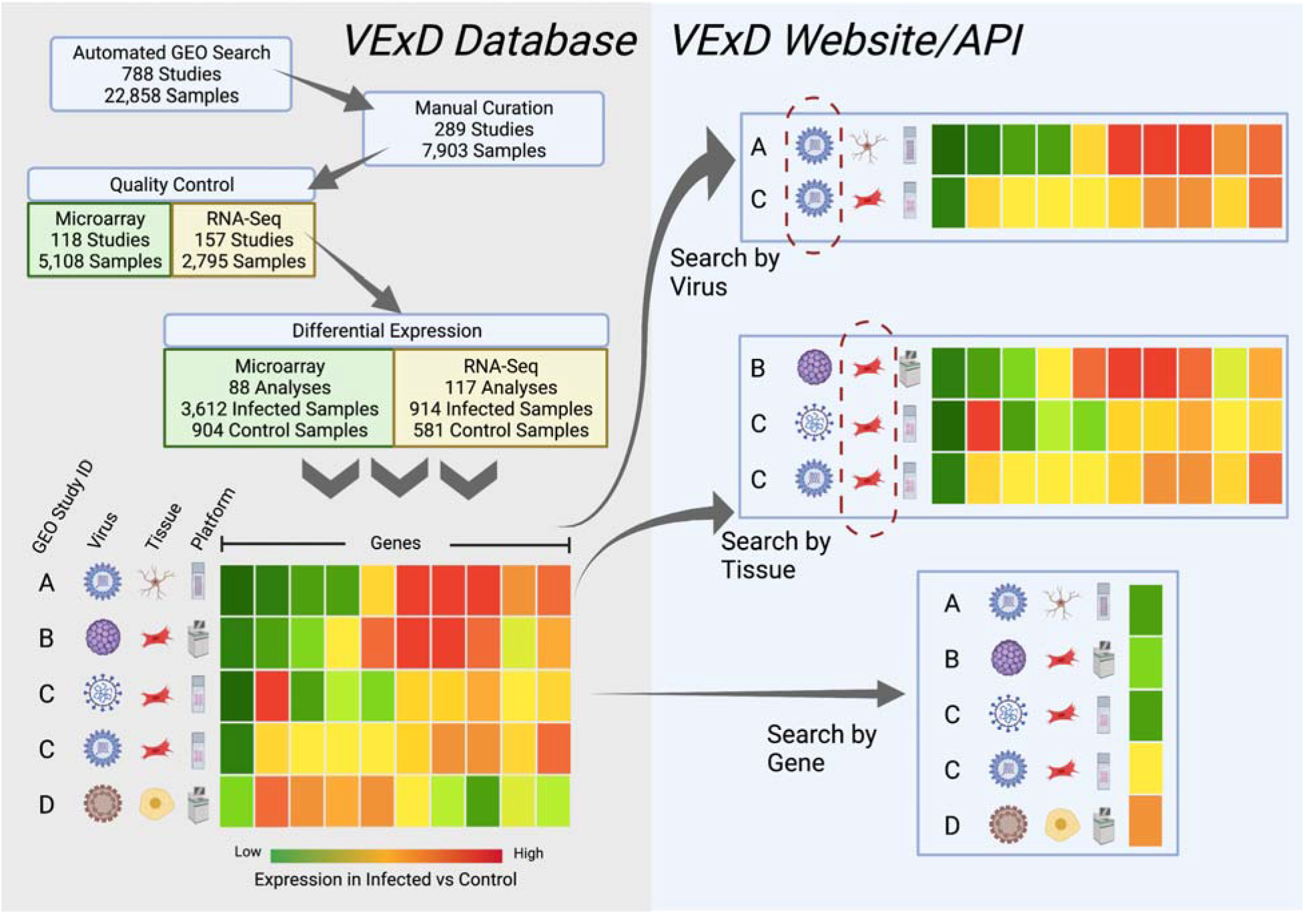
Overview of the VExD methods and website. The underlying database (left side) contains differential expression results for all viral infections of human cells or tissues found in GEO. Results are annotated with the source GEO ID, the virus species, the cell/tissue type (using the Brenda Tissue Ontology), and the experimental platform used in the original study. Users of the VExD website (right side) can search the results by virus or tissue type or can search by gene to see a global view of gene expression in response to viral infection. Static database dumps are available for download, and an extensive API is available for programmatic access.

## Results

### Identification of virus infection samples

We sought to identify and collate a comprehensive collection of experiments that show gene expression differences in human cells after infection with a virus. The Gene Expression Omnibus (GEO) is a large repository of gene expression experiments, making it a natural source for our collection efforts. Before identifying experiments in GEO that involve viral infection, we first assembled a comprehensive list of viruses known to infect humans (**Supplemental Table 1**, details in Methods). We then searched all GEO sample records for any virus name, alias, or abbreviation, and manually curated the results to our final set of 7,903 samples in 289 experiments of viral infection of human cells (further details in Methods). In addition to verifying that the samples underwent viral infection (or were a matched uninfected control), we standardized both the virus name and the cell or tissue type of the sample. Viruses were named according to the species name defined by the International Committee on Taxonomy of Viruses (Walker et al., 2020). For cell/tissue identification, samples were annotated with the most relevant term from the BRENDA Tissue Ontology (BTO) (Gremse et al., 2011).

### Uniform data processing and quality control

Because the data in question span approximately seventeen years of genomic technology and analysis improvements, we developed a uniform data processing and quality control pipeline. Using updated definition files from the BrainMap project (Dai et al., 2005), we discarded outdated microarray probes and used current definitions of genes across all samples for both microarray and RNA-Seq experiments. Briefly, RNA-Seq samples were quantified with kallisto (Bray et al., 2016) using Ensembl gene definitions and microarrays were processed with Robust Multi-chip Analysis (RMA) (Irizarry et al., 2003) using BrainMap custom CDF files that use the same gene definitions as the RNA-Seq samples. Following quality control (see Methods), 3.7% of microarray samples and 11% of RNA-Seq samples were removed. Differential analysis comparing gene expression levels between infected and uninfected cells within a study were performed using Welch’s T-test.

### The VExD web portal

The complete results of these curation and uniform processing efforts constitute VExD, the Virus Expression Database, freely available at https://vexd.cchmc.org. VExD offers the ability to search for experiments by virus and/or tissue type and allows the download of differential expression results for each experiment. In addition, users can search for individual genes or gene sets to examine their relative expression levels across all experiments contained in VExD.

The Downloads page allows users to download the complete list of GEO experiments within VExD, the complete differential expression results across all experiments, or differential expression results for each virus. Finally, computational users can make use of the Application Programming Interface (API) of VExD to perform any search available on the web site from the command line or their own programming environment.

### Validation

We sought to validate the VExD data processing methodology. We first examined the behavior of the 98 core interferon-stimulated genes (ISGs) that Shaw and colleagues (Shaw et al., 2017) identified as conserved across vertebrates **(Figure 2)**. Since ISGs are broadly activated in response to viral infection, we expected to see these genes largely upregulated in VExD. Indeed, we find that the stochastic superiority statistic for ISGs – that is, the probability that a randomly selected ISG measurement is larger than a randomly selected non-ISG measurement – is 64.7%, compared to a random expected value of 50%. Further, ISG fold changes are significantly distinct from all other (non-ISG) genes (p-value < 10^−30^, two-sided Brunner-Munzel test). Conversely, random selection of 98 genes shows no substantial change **(Supplemental Figure 1:** stochastic superiority statistic of 49.6%, p-value=0.17), as expected. The VExD website includes functionality (https://vexd.cchmc.org/enrich) to test input gene sets for enrichment.

**Figure 2.**
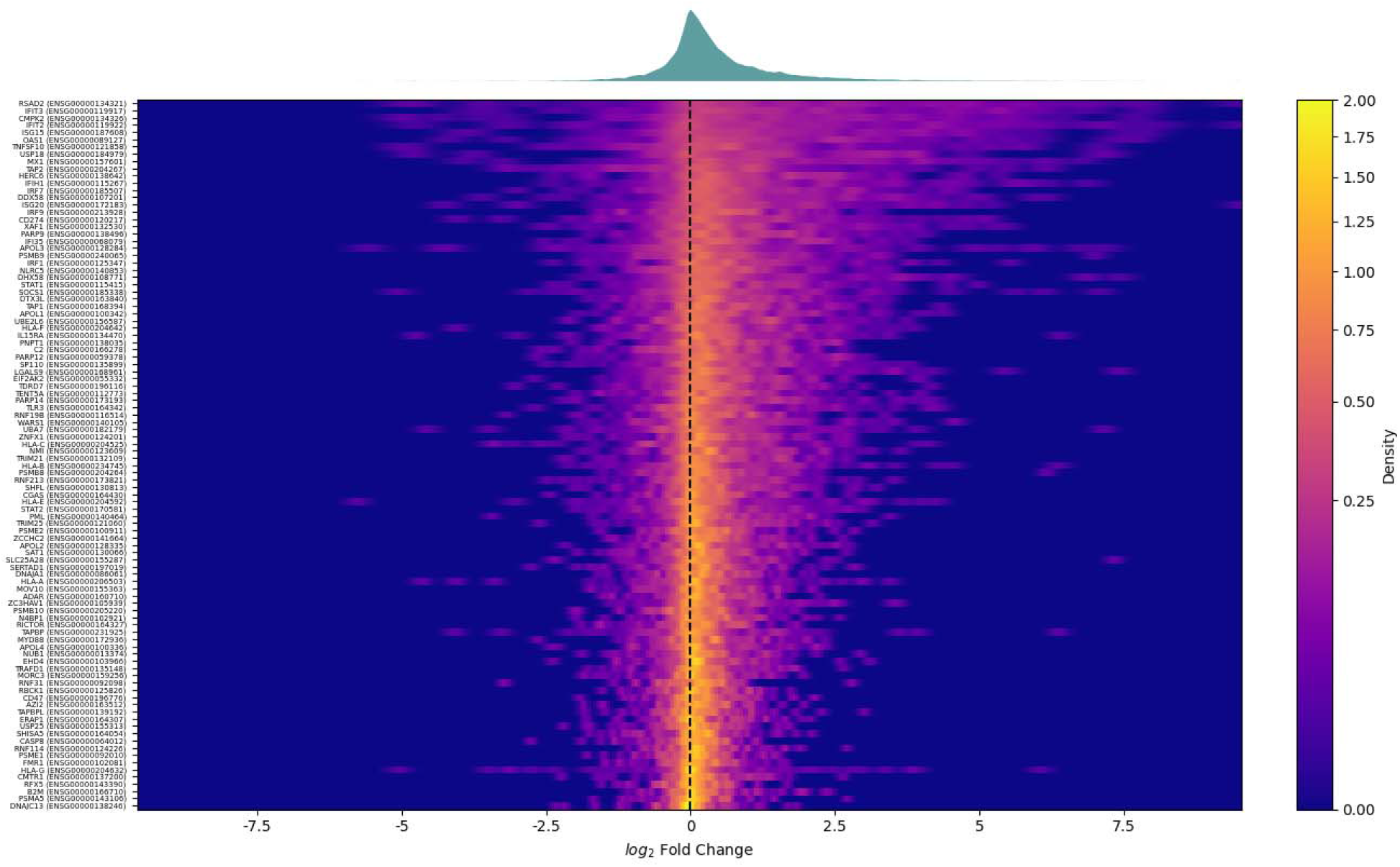
Pan-VExD expression of the core interferon-stimulated genes (ISGs) identified in (Shaw et al., 2017). Each row of the heatmap represents all differential expression results in VExD for a single gene, with colors indicating the density estimate of observed fold changes for that gene. The x-axis shows the log^2^ transformed fold change of infected samples versus their matched controls. A density plot of all 98 genes is shown above the heatmap. Note that both the heatmap and density plot have greater signal on the right side, indicating a global increase in gene expression, as expected for ISGs. Randomly selected genes do not produce this shift **(Supplemental Figure 1)**.

We next examined the ability of VExD data to highlight virus-specific effects. Tetherin is an ISG that physically binds virions to the host cell, preventing their export (Neil et al., 2008). It has been confirmed to be effective against a wide variety of enveloped viruses, including retroviruses (Neil et al., 2008), rhabdoviruses (Weidner et al., 2010), and filoviruses (Kaletsky et al., 2009). Indeed, when we examine the expression of tetherin in the experiments contained in VExD (Figure 3), we see that it is more likely to be overexpressed in response to infection with an enveloped virus than a virus with no envelope. The stochastic superiority statistic for tetherin expression after infection with an enveloped virus is 66.6%, with a Brunner-Munzel p-value of 5×10^−10^. Conversely, after infection with a naked (non-enveloped) virus, the stochastic superiority statistic is only 61.8%, with a p-value of 0.04. *Strikingly*, tetherin expression is uniformly upregulated in all 11 experiments of *Rhinovirus A* infection in VExD, even though rhinovirus does not have an envelope. When considering only the non-rhinovirus naked viruses, the stochastic superiority statistic falls to 49.5%, with a p-value of 0.9.

**Figure 3.**
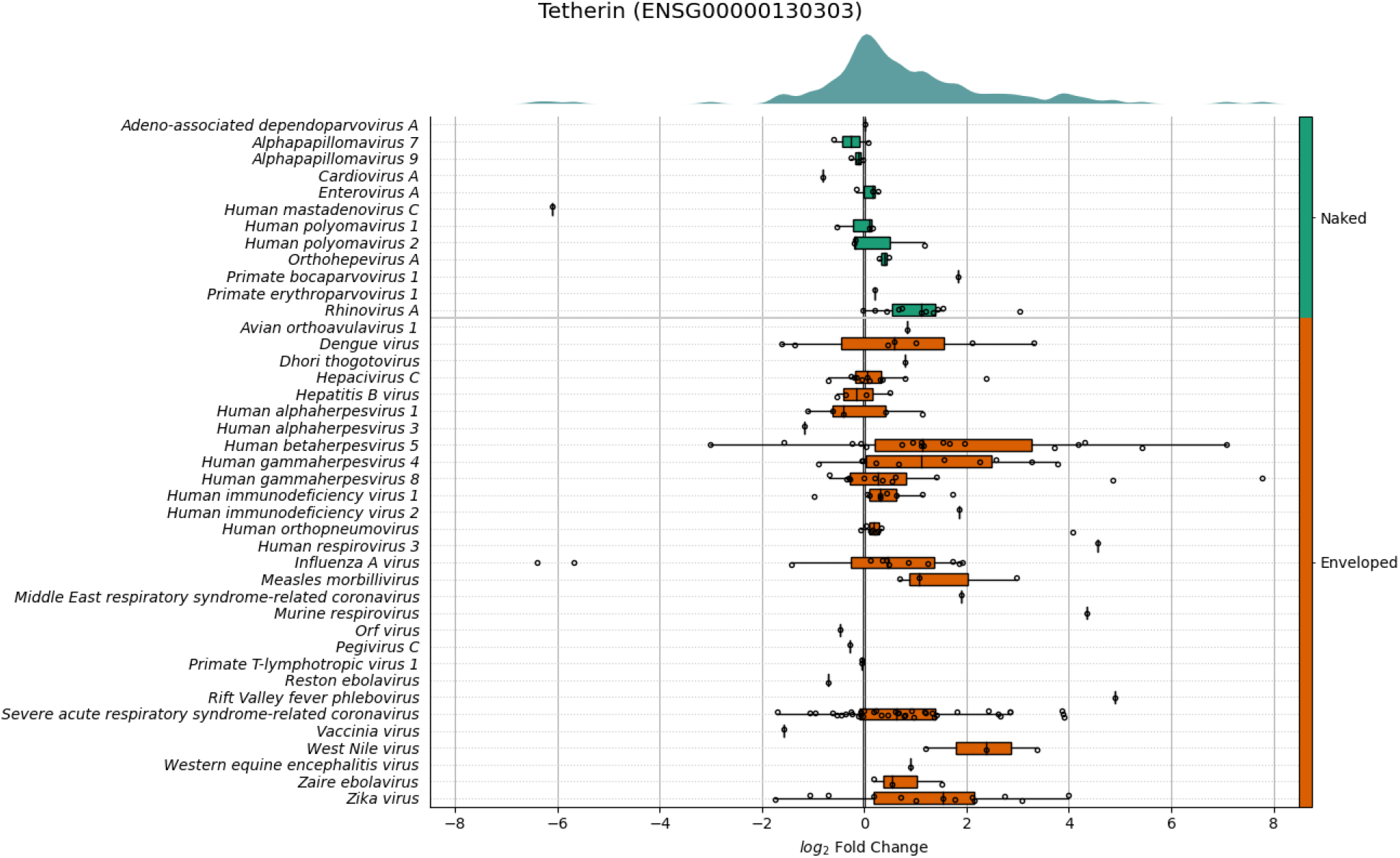
Expression of Tetherin (*BST2*) across all experiments in VExD. Experiments are stratified by virus (rows), which are then ordered by whether the virion has an envelope or not (right side). Within each virus, the log^2^ fold change of Tetherin in each distinct experiment is plotted as a dot. Box-and-whisker overlays are included as well. Broadly, Tetherin is overexpressed in response to enveloped viruses but is not overexpressed for naked (unenveloped) viruses, with the (novel) exception of Rhinoviruses.

We next examined the consistency of VExD results across experiments. A recent study (C. Wang et al., 2019) performed a time series of Epstein-Barr virus (*Human gammaherpesvirus 4*) infections against primary human B-cells. As such, this experiment is included in VExD, along with three prior experiments that also infected B-cells with EBV. Notably, Wang *et* al used RNA-seq to assess gene expression differences, while the other three experiments in VExD used microarrays. When we assess the differentially expressed genes highlighted in Figure 1 of Wang *et* al, we see that the direction of regulation is maintained in the other EBV-infected B-cell experiments contained in VExD **(Figure 4)**. These results are consistent with previous observations of the concordance between RNA-seq and microarray technologies (Rao et al., 2019; C. Wang et al., 2014), although the experiments in this case were performed in different laboratories with different experimental protocols.

**Figure 4.**
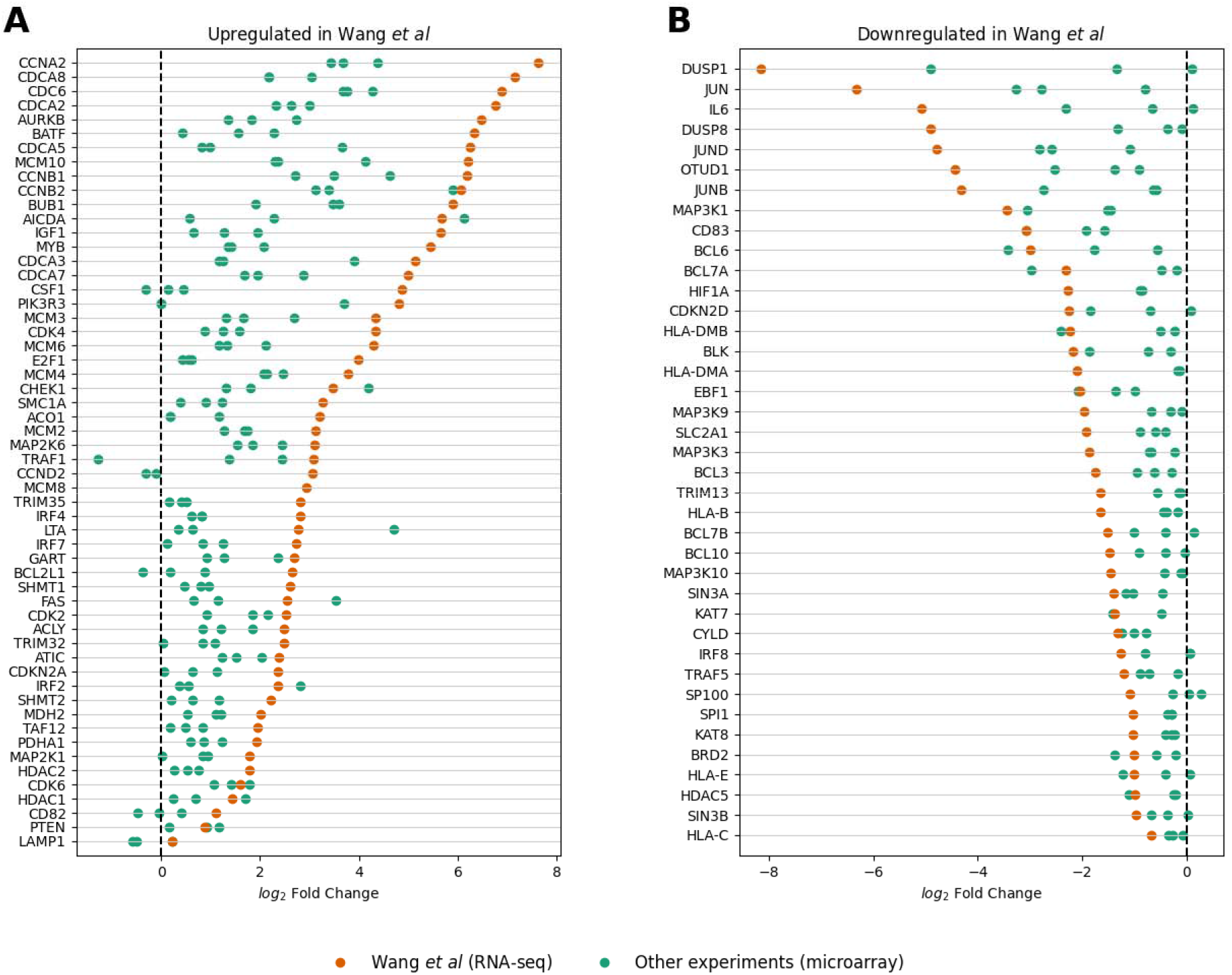
Differentially expressed genes identified by (C. Wang et al., 2019) show consistent expression across all similar experiments in VExD. In each panel, the genes identified by Wang et al are displayed in rows. Each dot represents the fold change of that gene in a single VExD experiment, after restricting to experiments of Epstein-Barr virus infection in human B-cells. Results from Wang *et al* are highlighted. Panel **A)** genes found to be upregulated in Wang *et al*, **B)** genes found to be downregulated in Wang *et al*.

Finally, we used VExD to add context to findings obtained from a model organism. A recent publication (Ma et al., 2021) demonstrated that the m^6^A reader gene *Ythdf2* is overexpressed in mouse natural killer cells after infection with murine cytomegalovirus (*Murid betaherpesvirus 1*). Using VExD, we find that the human ortholog YTHDF2 is also overexpressed in response to human cytomegalovirus (*Human betaherpesvirus 5*) infections, but not to other virus types **(Figure 5)**. In addition, although none of the VExD experiments were performed in natural killer cells, a tissue-specific response is apparent, with only fibroblasts and epithelium showing upregulation among the cell types assayed **(Supplemental Table 2)**.

**Figure 5.**
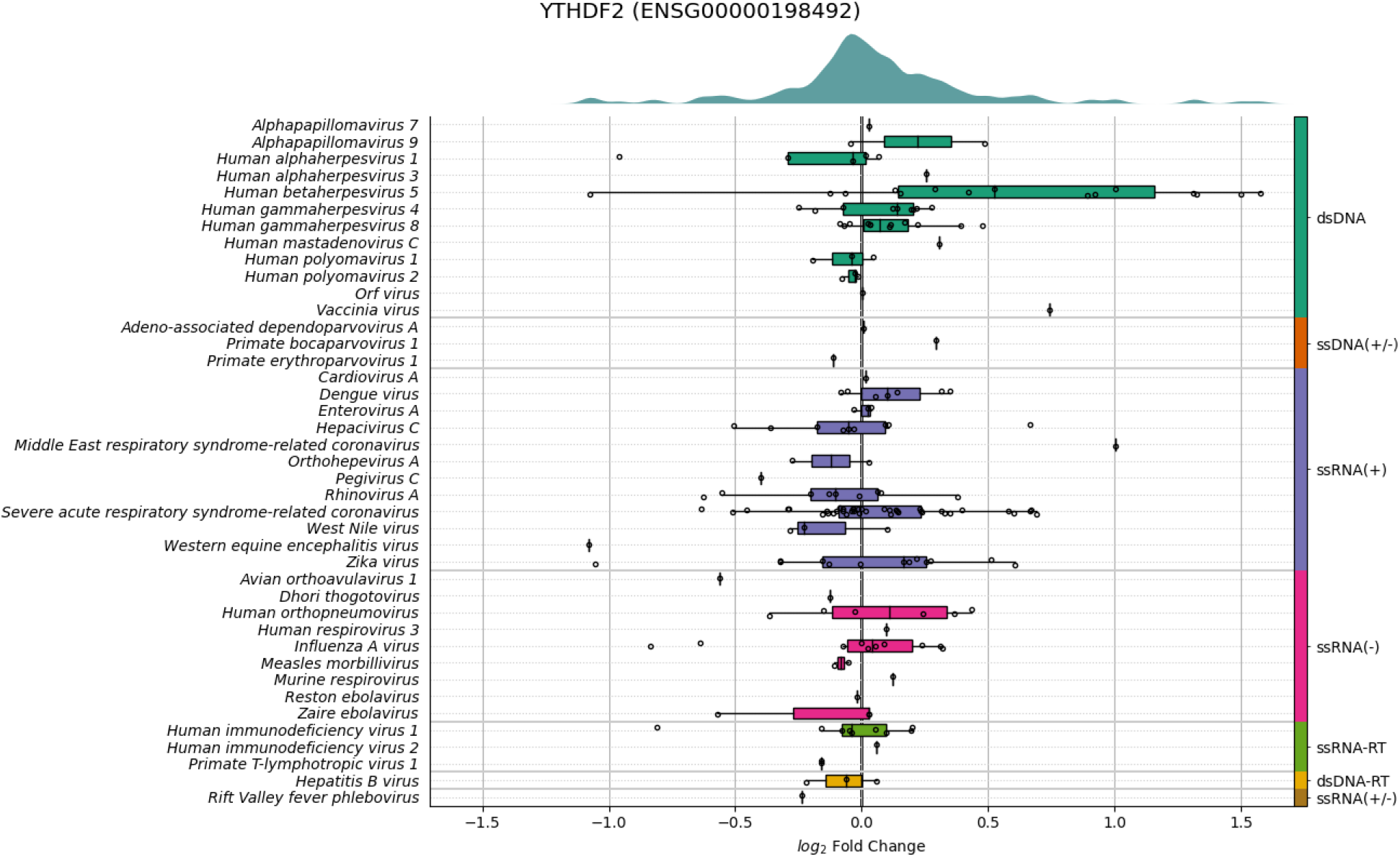
Expression of *YTHDF2* across all VExD experiments. Experiments are stratified by virus (rows), which are then ordered by genome type (right side). Within each virus, the log^2^ fold change of *YTHDF2* in each distinct experiment is plotted as a dot. Box-and-whisker overlays are also included. Note that *YTHDF2* is strongly upregulated in response to *Human betaherpesvirus 5* (CMV) infections, but not other viruses.

## Discussion

We present the Virus Expression Database (VExD), a comprehensive, curated resource of transcriptomic studies of viral infection in human cells. VExD contains 7,903 samples from 289 experiments, all processed through uniform quality control, quantification, and differential expression pipelines. The VExD website (https://vexd.cchmc.org/) presents these results in a simple searchable form, as a bulk download, or via an Application Programming Interface. In addition, the VExD website allows users to query the response of genes to a viral infection, stratified by virus and tissue type through real-time data visualizations.

We demonstrate the utility of VExD using four case studies. In the first, we show that the interferon response is highly consistent across viruses, as expected **(Figure 2)**. In the second **(Figure 3)**, we demonstrate the ability of VExD to highlight virus-specific effects by showing that the gene expressing tetherin is specifically upregulated in enveloped viruses. In our third case study **(Figure 4)**, we show that the results in VExD are consistent across experiments, laboratories, and analytic platforms. Specifically, the genes identified by Wang et al (C. Wang *et al*., 2019) as significantly regulated by EBV in B cells show similar patterns in three independent experiments in VExD, despite the Wang group using RNA-seq and the other experiments using microarray. In our final case study **(Figure 5)**, we examine a report of a novel antiviral gene (*Ythdf2*) found in a model organism (mouse). Using the data contained in VExD, we find that the human ortholog is also upregulated in humans in a virus- and cell-type specific manner. In this manner, the breadth of experimental data captured by VExD provides a broader context to investigators, allowing for more nuanced hypothesis generation.

VExD is a novel and extensive resource that will be of broad benefit to the community. However, it does have some limitations. First, VExD is limited to the studies that have been deposited in GEO, which means that particular virus or cell types may be underrepresented. In addition, although we performed extensive quality control of the transcriptomic assays deposited in GEO before including them **(Methods)**, we could not verify all experimental details, such as whether the appropriate multiplicity of infection or cell type were assayed. As such, it is possible that VExD contains experiments that do not represent successful infections. Second, VExD relies upon manual curation of automatically generated GEO search results. While this is a labor-intensive process that is susceptible to human error, in our experience a fully automated process does not adequately capture the complex descriptions contained in GEO.

To our knowledge, VExD is the largest available resource of gene expression data following viral infection. Using the website or the API, investigators can use VExD to identify relevant transcriptomic studies, query gene expression levels in a system of interest, or investigate the relevance of individual genes or gene sets in the antiviral response. As new virus gene expression data continue to become available, we anticipate that future updates to VExD will continue to improve its utility through the incorporation of these additional datasets.

## Materials & Methods

### Identification of samples

We assembled a list of viruses known to infect humans based on annotations in ViralZone (Hulo et al., 2011), UniProt (The UniProt Consortium, 2019), the Virus-Host DB (Mihara et al., 2016), and literature curation studies (Taylor et al., 2001; Woolhouse et al., 2012), as well as common synonyms and abbreviations. Following the taxonomy published by the International Committee on Taxonomy of Viruses (Walker et al., 2020), we standardized all viruses to the level of species. Our final list **(Supplemental Table 1)** comprises 332 human-infecting species.

Using a tool we developed for a previous publication (Harley et al., 2018), we searched GEO for any studies mentioning a virus name or abbreviation in the study or sample description. Only samples annotated as human were considered. To enable standardized processing, we also required that studies use either the Affymetrix microarray or Illumina bulk RNA sequencing platforms. The 22,858 samples identified with the automated scanner were then manually curated to remove false positive results (**Figure 1**, top left). Samples were removed for a variety of reasons, including: 1) wrong platform, 2) use of a virus as a molecular biology tool rather than an infection (e.g., to immortalize cells or as a gene promoter), or 3) the virus name abbreviation identified by the scanner was not used in a virology context (e.g., Measles Virus = MV = Megavolts). The remaining 7,903 samples were annotated with Brenda Tissue Ontology (BTO) terms representing the assayed cell type or tissue (Gremse et al., 2011) and were loaded into a Mongo database.

### Data Quality Control

RNA-Seq samples were quantified using kallisto (Bray et al., 2016) with the --bias parameter to account for sequence-specific biases. For paired-end samples, kallisto was allowed to derive the fragment length distribution from the data (default behavior). For single-end samples, the distribution was specified as having a mean length of 300nt and a standard deviation of 30nt (parameters “-l 300 -s 30”). For all samples, the reference transcriptome was the Ensembl human gene set (version 102). Final gene-level quantities were obtained by summing the TPM for all associated transcripts. Samples with too few usable reads (pseudo-aligned reads < 1.5 million) or poor overall alignment (< 18% reads pseudo-aligned) were removed.

For microarray studies, BrainMap version 25 CDFs (Dai et al., 2005) that remap probes onto Ensembl v102 genes were used. All microarrays of a single array type were analyzed together using Robust Multi-chip Analysis (RMA) (Irizarry et al., 2003) as implemented in the Affymetrix Power Tools. Samples were removed if the residual error in the RMA model was too large (mad_residual_mean > 0.80) or if the overall signal was too low (pm_mean < 65). For RNA-seq and microarray platforms, studies with fewer than 60% of their samples passing these criteria were removed. In total, 234 of 2,129 (11%) RNA-seq samples and 186 of 4,964 (3.7%) microarray samples were removed by these quality control steps.

### Differential Expression Quantification

VExD contains differential expression results between comparable virus infected samples and uninfected controls, allowing users to easily determine the human gene expression changes observed for each virus. Comparable samples are defined as those coming from the same study, using the same tissue or cell type, and the same technological platform, with at least two infected samples and two uninfected control samples. A total of 205 differential expression analyses involving 4,526 infected samples and 1,485 uninfected controls are available in VExD **(Figure 1)**. All tests used a two-sided Welch’s T test with unequal variance and Benjamini-Hochberg correction.

### Gene Enrichment Analysis

When analyzing gene sets, we compare the fold changes observed for the genes in the set across all experiments in VExD to all other gene observations. Specifically, we employ the Brunner-Munzel test (Brunner & Munzel, 2000; Neubert & Brunner, 2007) to determine whether the distribution of values associated with the gene set are significantly different from the remaining values in VExD. The Brunner-Munzel test is similar to the Wilcoxon-Mann-Whitney test but does not assume that the variance within each population is the same.

The stochastic superiority statistic is called the “common language effect size” in some sources, and is calculated as described in (Brunner et al., 2018), where it is named 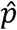.

### VExD Website and API

The VExD website was created with the Python Flask framework and is served using the Apache webserver with the mod_wsgi module. Users can search by any combination of virus and tissue type to view overall gene expression levels, or search for individual genes across all studies by symbol, alias, or Ensembl ID **(Figure 1)**. Results pages display the virus, cell type, fold change, and adjusted p-value for a given gene in all studies. Download links provide study and differential expression information in both a tab-delimited text file and a compressed, machine-readable format (Apache Parquet). VExD also supports an Application Programming Interface (API) that can be used to programmatically query the database on the command-line.

### Data/Source Code Availability

The VExD website (https://vexd.cchmc.org) is freely available, with no registration required. Differential expression results can be downloaded from VExD on a per-study or per-virus basis, or all in a single file. Data curation results are also available for download. Figures 2 and S1 in this paper can be regenerated from the “Enrichment” tab of the website. Source code to generate Figures 3 and 4 is included as supplemental material. Figure 5 can be regenerated from the “Genes” tab of the website. Source code for the website is available at https://github.com/pdexheimer/vexd.

The genes used in Figure 4A were the “Genes of interest” from Groups 1, 2, and 3 (upregulated upon infection) from Figure 1 of (C. Wang et al., 2019). The genes used in Figure 4B were similarly chosen from Groups 7 and 8 of the same figure. Groups 4, 5, and 6 showed more complicated, time-dependent expression patterns, and were thus not relevant for a simple “infected versus control” comparison. Gene lists used in all figures are available as **Supplemental Table 3**.

## Acknowledgements

We thank Leah Kottyan and Erica DePasquale for helpful feedback on the manuscript. We thank Carmy Forney and Leah Kottyan for assistance with manual dataset annotations. We thank Kevin Ernst for technical assistance with web site development. We thank members of the Weirauch and Kottyan lab for feedback on the VExD web site. This work was funded by National Institutes of Health grants R01NS099068, R01HG010730, R01GM055479, U01AI130830, R01AI024717, U01AI150748, R01AI148276, and P01AI150585 to M.T.W.

## Author contributions

M.T.W., P.J.D., and K.R came up with the concept for the project. P.J.D. performed all computational analyses and wrote most of the manuscript. M.T.W. and K.R. supervised the project. M.P. wrote the computational code for querying the GEO database. All authors contributed to the writing and editing of the manuscript and approve of the contents of the manuscript.

---

**Supplemental Figure 1.**
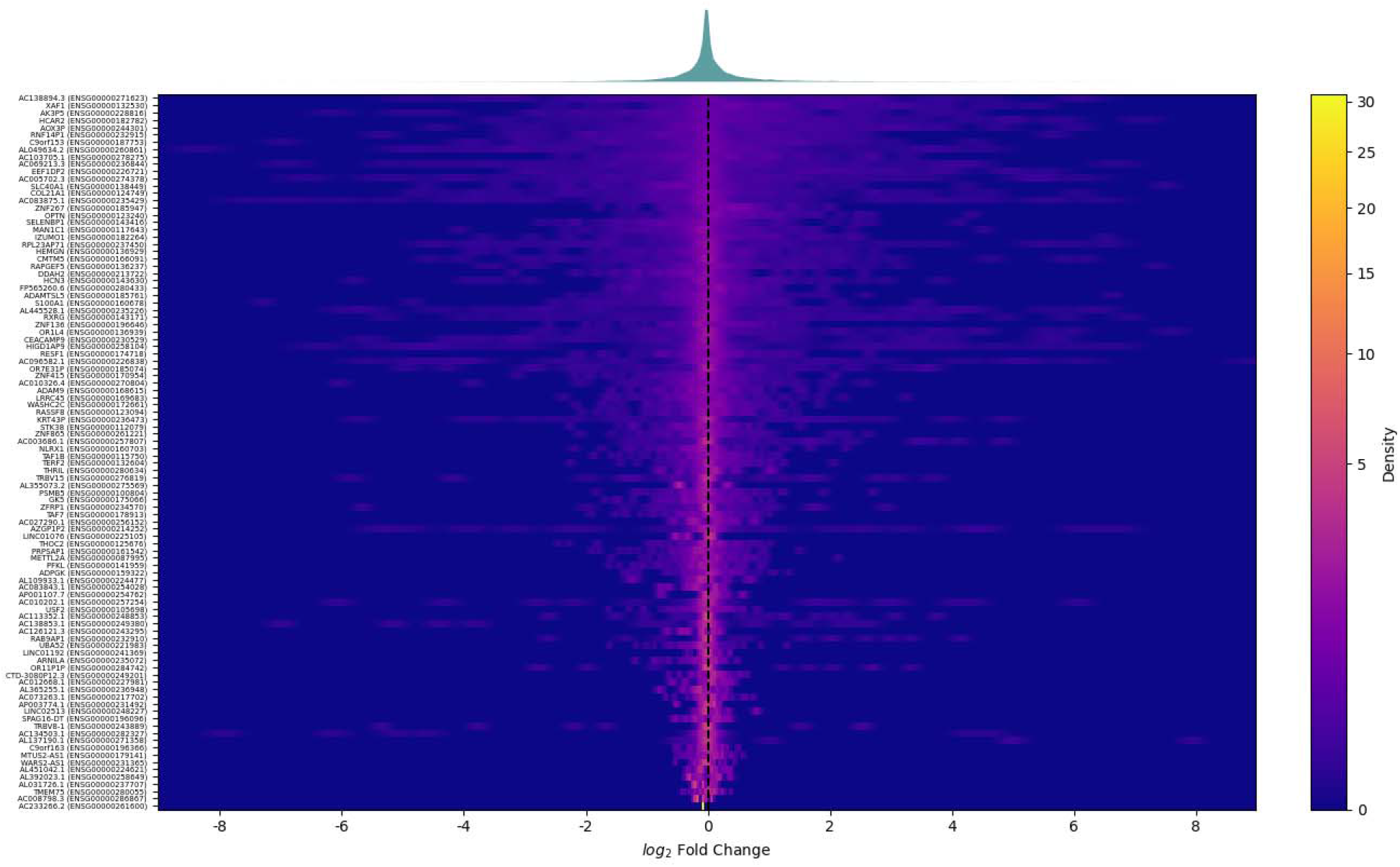
Pan-VExD expression of 98 randomly-selected genes. As in Figure 2, each row represents the fold change of a single gene across all experiments within VExD, and the top panel contains a density plot of all 98 genes. As expected, these genes are largely unchanged in response to viral infection. The VExD website includes functionality to regenerate this figure with different random genes or user-selected gene sets.

**Supplemental Table 1 (Separate file) –** The species name, aliases, and abbreviations for all human-infecting viruses eligible for inclusion in VExD.

**Supplemental Table 2** (Separate file) – Per-experiment details for *YTHDF2* expression following cytomegalovirus (*Human betaherpesvirus 5*) infection, highlighting the cell type-specific nature of this gene’s regulation. This table is a subset of the information available from https://vexd.cchmc.org/gene?q=ENSG00000198492 or the results endpoint of the VExD API.

**Supplemental Table 3** (Separate file) – The genes used in Figures 2, 4, and S1 of this paper.

**Supplemental File 4** (Separate file) – the Python source code used to generate Figure 3 of this paper.

**Supplemental File 5** (Separate file) – the Python source code used to generate Figure 4 of this paper.

## Notes

### Competing Interest Statement

The authors have declared no competing interest.

### Summary of Updates

Expanded introduction, results, and discussion; added case studies; added Figures 2-5; added functionality to website; added supplemental files

