## Supplemental Files 4 & 5 for "VExD: A curated resource for human gene expression alterations following viral infection": File S4.py

```
import requests
import pandas as pd
import matplotlib as mpl
mpl.use('agg')
from matplotlib.cm import ScalarMappable
from matplotlib.colors import BoundaryNorm
from matplotlib.figure import Figure
import numpy as np
from scipy.stats import gaussian_kde

ensembl_host = 'https://rest.ensembl.org'
vexd_host = 'https://vexd.cchmc.org'

def get_ensembl_id(gene):
    """
    Given a (human) gene symbol, queries the Ensembl REST API to determine
    the Ensembl gene ID (ENSGXXXXX) for that gene.

    If the query returns multiple Ensembl IDs, only the first is returned
    by this function. If the query returns no results, this function returns
    None

    Raises an Exception if the REST call errors
    """
    r = requests.get(
        f'{ensembl_host}/xrefs/symbol/homo_sapiens/{gene}',
        {
            'content-type': 'application/json',
            'external_db': 'HGNC'
        }
    )
    r.raise_for_status()
    result = r.json()
    for ens_gene in result:
        return ens_gene['id']

    return None

def get_vexd_gene_results(ensembl_id):
    """
    Gets the differential expression results from VExD for a single gene.
    Raises an Exception if the HTTP query fails
    """
    r = requests.post(
        f'{vexd_host}/api/v1/results',
        json={'gene': ensembl_id}
    )
    r.raise_for_status()
    return r.json()

def get_vexd_virus_info(species):
    r = requests.post(
        f'{vexd_host}/api/v1/virus',
        json={'name': species}
    )
    r.raise_for_status()
    return r.json()

def get_envelope_status(df):
    """
    Given a DataFrame containing a "virus" column, returns a boolean
    Series that indicates whether that virus has an envelope.

    Designed to be used in a DataFrame.assign() call
    """
    annotation = pd.DataFrame({'virus': df['virus'], 'envelope': False})\
        .drop_duplicates(ignore_index=True)\
        .set_index('virus')
    for virus in annotation.index.to_list():
        info = get_vexd_virus_info(virus)
        annotation.loc[virus, 'envelope'] = info['envelope']
    return df.join(annotation, on='virus', sort=False)['envelope']


def _get_class_boundaries(data, category):
    """
    Internal function to find the boundaries between categories (specified 
    as a column name) in the data
    """
    results = []
    viruses = data.drop_duplicates('virus', ignore_index=True)
    cat_col = viruses.columns.get_loc(category)
    for i in range(1, viruses.shape[0]):
        if viruses.iat[i, cat_col] != viruses.iat[i-1, cat_col]:
            results.append(i+0.5)
    return results

def add_colorbar(figure, axes, boundaries, names):
    """
    Creates a colorbar that corresponds to categories in the input,
    adds it to the figure.
    """
    augmented_bounds = [axes.get_ylim()[1],] + boundaries + [axes.get_ylim()[0],]
    cbar = figure.colorbar(
        ScalarMappable(
            BoundaryNorm(augmented_bounds, len(augmented_bounds)-1), 
            mpl.colormaps['Dark2']
        ),
        spacing='proportional',
        ticks=[],
        fraction=0.02,
        aspect=50,
        pad=0,
        ax=axes
    )
    cbar.ax.invert_yaxis()
    cbar.set_ticks(
        pd.Series(augmented_bounds).rolling(2).mean().tail(-1).to_list(),
        labels=names,
        minor=True
    )

ensembl_id = get_ensembl_id('tetherin')
results = pd.DataFrame(get_vexd_gene_results(ensembl_id))\
    .assign(envelope=get_envelope_status)\
    .sort_values(by=['envelope', 'virus'])
labels = []
data = []
for name, subset in results.groupby('virus', sort=False):
    labels.append(name)
    data.append(subset['logfc'].to_list())

# Main figure creation and plotting
fig = Figure(figsize=(13, 8), layout="constrained")
ax = fig.add_subplot(10, 1, (2,10)) # Bottom 90%
bplot = ax.boxplot(
    data,
    labels=labels,
    vert=False,
    patch_artist=True,
    widths=0.7,
    medianprops={'color': 'black'},
    showfliers=False,
    showcaps=False,
)
# Color the boxplots by genome composition
for patch, genome_idx in zip(bplot['boxes'], results.drop_duplicates('virus').groupby('envelope', sort=False).ngroup()):
    patch.set_facecolor(mpl.colormaps['Dark2'](genome_idx))
# Plot the individual points over the boxplots, with a small amount of jitter
for y, x in enumerate(data):
    ax.plot(
        x, 
        np.random.uniform(-0.2, 0.2, len(x))+y+1, 
        linestyle="none", 
        color='black', 
        markersize=3, 
        marker='o', 
        fillstyle='none',
    )
# Set up the axes and labels
ax.set_xlabel("$log_2$ Fold Change")
ax.set_ylabel("")
ax.set_yticklabels(
    labels, 
    fontdict={'fontstyle': 'italic'}
)
max_extent = max([abs(x) for x in ax.get_xlim()])
ax.set_xlim(-max_extent, max_extent)
ax.spines.top.set_visible(False)
ax.spines.right.set_visible(False)
ax.invert_yaxis()
# Hide the tick marks on the y axis
for tick in ax.yaxis.get_major_ticks():
    tick.tick1line.set_markersize(0)
    tick.tick2line.set_markersize(0)
# Add grid lines and the lines highlighting the 0 point and the change in genome composition
ax.axvline(x=0, color='k', linestyle='-', linewidth=2, zorder=0)
class_bounds = _get_class_boundaries(results, 'envelope')
for bnd in class_bounds:
    ax.axhline(y=bnd, color='0.8', linestyle='-')
ax.xaxis.grid(True)
ax.grid(axis='y', linestyle=':', color='0.8')
# Title and colorbar
fig.suptitle(f"Tetherin ({ensembl_id})", fontsize="x-large")
add_colorbar(fig, ax, class_bounds, ['Naked', 'Enveloped'])
# Density plot of all points
upper_ax = fig.add_subplot(10, 1, 1, sharex=ax, frame_on=False)
density = gaussian_kde(results['logfc'].to_numpy())
density.set_bandwidth(density.factor / 4.)
density_x = np.linspace(ax.get_xlim()[0], ax.get_xlim()[1], 200)
upper_ax.fill_between(density_x, density(density_x), facecolor='cadetblue')
upper_ax.set_axis_off()
# Finally, write out the PNG
fig.savefig('Figure 3.png')
```
