## Supplemental Files 4 & 5 for "VExD: A curated resource for human gene expression alterations following viral infection": File S5.py

```
# Additional libraries that need to be accessible:
#  * pyarrow (used by pandas.read_parquet)
#  * openpyxl (used by pandas.read_excel)
import pandas as pd
import matplotlib as mpl
mpl.use('Agg')
from matplotlib.figure import Figure
from matplotlib.lines import Line2D

ebv_results = pd.read_parquet('https://vexd.cchmc.org/download/Human_gammaherpesvirus_4.parquet')
bcells = ebv_results.\
    xs('B-lymphocyte', level='bto_name', axis='columns').\
    xs('log_fc', level='data', axis='columns')
bcells.columns = bcells.columns.droplevel(['virus', 'bto_id', 'platform'])
bcells.reset_index('ensembl_id', drop=True, inplace=True)

genes = pd.read_excel('Table S3.xlsx', sheet_name='Figure 4')
up = genes['Up-regulated'].rename("genes").dropna()
down = genes['Down-regulated'].rename('genes').dropna()

def plot_genes(ax, data, highlight=None):
    if highlight is not None:
        data = data.sort_values(by=[highlight], key=lambda x: x.abs(), ascending=False)
    ax.set_yticks(
        list(range(data.shape[0])),
        reversed(data.index.values)
    )
    for series in data.columns.values:
        if series != highlight:
            ax.scatter(data[series], list(reversed(range(data.shape[0]))), color=mpl.colormaps['Dark2'](0))
    if highlight is not None:
        ax.scatter(data[highlight], list(reversed(range(data.shape[0]))), color=mpl.colormaps['Dark2'](1))
    ax.axvline(x=0, color='black', linestyle='dashed')
    ax.grid(axis='y', color='#CCC')
    ax.set_xlabel(r'$log_2$ Fold Change')
    ax.set_ylim(bottom=-1, top=data.shape[0])

fig = Figure(figsize=(13, 10), layout='constrained')
fig.get_layout_engine().set(w_pad=0.5, h_pad=0.2)
subfigs = fig.subfigures(nrows=2, height_ratios=[12,1], hspace=0)
ax_a, ax_b = subfigs[0].subplots(ncols=2)
legend_ax = subfigs[1].subplots()
plot_genes(
    ax_a,
    pd.DataFrame(up).join(bcells, on="genes").set_index('genes'),
    highlight='GSE125974'
)
plot_genes(
    ax_b,
    pd.DataFrame(down).join(bcells, on="genes").set_index('genes'),
    highlight='GSE125974'
)
ax_a.text(-0.15, 1.02, 'A', transform=ax_a.transAxes, fontsize=24, fontweight='bold')
ax_a.set_title(r'Upregulated in Wang $et\ al$')
ax_b.text(-0.15, 1.02, 'B', transform=ax_b.transAxes, fontsize=24, fontweight='bold')
ax_b.set_title(r'Downregulated in Wang $et\ al$')
legend_ax.set_axis_off()
legend_ax.legend(
    [
        Line2D([], [], color=mpl.colormaps['Dark2'](1), marker='o', linestyle='None'),
        Line2D([], [], color=mpl.colormaps['Dark2'](0), marker='o', linestyle='None'),
    ],
    [
        r'Wang $et\ al$ (RNA-seq)',
        'Other experiments (microarray)'
    ],
    loc='center', 
    ncols=2,
    frameon=False,
    fontsize='large'
)
fig.savefig('Figure 4.png')
```
